# Analyses of the 5’ Ends of *Escherichia coli* ORFs

**DOI:** 10.1101/2021.08.27.457967

**Authors:** Michael Ward, James F. Curran

**Affiliations:** Department of Biology, Wake Forest University, PO Box 7325, Winston-Salem, NC 27109, USA

## Abstract

Sequence biases at 5’ ends of coding sequences differ from those of the remainder of ORFs, reflecting differences in function. Internal sequence biases promote translational efficiency by several mechanisms including correlating codon usage and tRNA concentration. However, the early region may also facilitate translational initiation, establishment of the reading frame, and polypeptide processing. Here we examine the beginnings of the ORFs of an *Escherichia coli* K12 reference genome. The results extend previous observations of A-richness to include an overabundance of the AAA triplet in all reading frames, consistent with the hypothesis that the beginnings of ORFs contribute to initiation site accessibility. Results are also consistent with the idea that the first two amino acids are under selection because they facilitate solvation of the amino-terminus at the end of the ribosomal exit channel. Moreover, serine is highly overrepresented as the second amino acid, possibly because it can facilitate removal of the terminal formylmethionine. Non-AUG initiation codons are known to be less efficient than AUG at directing initiation, presumably because of relatively weak base pairing to the initiator-tRNA. But non-UAG initiation codons are not followed by unusual 3’ nearest neighbor codons. Moreover, the four NUG initiation codons do not differ in their propensity to frameshift in an assay known to be sensitive to base pair strength. Altogether, these data suggest that the 5’ ends of ORFs are under selection for several functions, and that initiation codon identity may not be critical beyond its role in initiation.

## Introduction

The 5’ ends of bacterial coding sequences must achieve several goals. They contribute to translational initiation and establish the proper reading frame, and the resultant peptides must initiate polypeptide passage through the ribosomal exit channel and facilitate formylmethionine (f-Met) removal. These properties do not apply to internal sites, and one might therefore expect that sequence biases at the beginnings of ORFS would differ from other gene sequence biases.

Gene sequences upstream of translational initiation sites have very well-characterized features such as Shine-Dalgarno elements^1^, an initiation codon from a restricted set of triplets, and an appropriately sized spacer between them^2^. In addition, initiation regions tend to be adenine- and uridine-rich^3^. It was predicted that these biases would facilitate translation by reducing the propensities for secondary structures that could otherwise interfere with translation initiation by blocking ribosomal access. Moreover, in at least some cases, ribosomes may bind further upstream to facilitate initiation by increasing the local ribosomal concentration^4^.

In addition, message sequences internal to open reading frames (ORFs) show extensive and highly significant non-randomness in codon usage^5^. It is thought that those internal biases contribute to translational efficiency. For example, codon bias generally matches codon frequencies to tRNA abundance, which should promote the speed and accuracy of expression. In many cases codons read by the same tRNA can be translated at different rates, and highly expressed genes tend to use the more rapidly translated codons^6,7^. Additionally, sites that are frameshift-prone tend to be avoided in genes^8,9^. Codon contexts are also nonrandom; such patterns may be due to effects of neighboring message nucleotides on codon recognition, or on interactions between peptidyl and aminoacyl tRNAs in the ribosomal A and P sites^10,11^. Moreover, ORFs typically exhibit a triplet “phase bias” such as the “G-nonG-N’’ pattern found for a subset of *E. coli* genes, which may be due to demands on amino acid usage in proteins^8^.

The sequence biases observed at the beginnings of ORFs are very different from those observed internally^2^. For example, the initiation codon is usually AUG, which is otherwise not common. In addition, the entire initiation region is A- and U-rich. This apparent selection for initiation site availability must affect amino acid usage, and such effects may have significance for protein function. This work explores sequence bias at the beginning of the average *E. coli* ORF. We study the nucleotide, codon and encoded amino acid usage for the region just 3’ of the initiation codon to determine the probable functional significance of these patterns.

## Materials and Methods

### Sequence analyses

We studied the *E. coli* K-12 substrain MG1655 complete genome sequence Genbank (accession U00096, Blattner et al. 1997; September 24, 2018 draft)^12^. All programs were written in Python version 3, running on the Google Colaboratory cloud computing platform. Programs are available at: https://github.com/Curran-WFU.

### Analysis of ribosomal frameshifting

Frameshifting was studied *in vivo* using a plasmid-borne *lacZ* reporter system that has been described^13^. Briefly, the *E. coli* K12 strain MY600 [Δ(*pro lac*) *ara, thi*] carries a *lacZ* deletion, and the plasmid carries the only functional *lacZ* allele. The plasmid that was described, pJC1168, was modified to contain a HindIII site in the region between the Shine/Dalgarno element and the initiation codon. Experimental plasmid constructs were made by cloning 30-nucleotide, double-stranded oligonucleotides between HindIII and BamHI sites. These constructs replace the initiation region with synthetic sites. Four pairs of sites were constructed that differ in their initiation codons (ATG, GTG, CTG and TTG). One member of each pair required frameshifting for *lacZ* expression while the other served as a non-frameshifting control.

## Results and Discussion

Previous work has shown that nucleotide and codon biases at the start of coding sequences differ from the patterns beyond initialization sites. However, the significance of these differences is not fully known. We wished to return to this topic using a complete genome sequence and address several questions. How far does the bias at initiation sites extend before it transitions to internal bias? What are the likely causes of initiation region bias? Does bias at initiation sites differ among genes that use AUG vs. other initiation codons? Does initiation codon identity affect the establishment of the correct reading frame? Our first task was to extract information on protein coding genes from an *E. coli* reference genome. We then address each question in turn.

### Parsing the *Escherichia coli* K-12 genome

We used the *E. coli* K-12 substrain MG1655 complete genome sequence. This genome has served as a standard *E. coli* reference for molecular biological research and has been refined extensively over the past two decades (originally published by Blattner et al. 1997)^12^. The number of annotated genes in the file is 4546. We analyzed only those that have open reading frames (ORFs) that start with known initiation codons (ATG, GTG, CTG, TTG, ATT, ATA, AAA) and end with standard termination codons (TAA, TAG, TGA). These criteria removed genes for functional RNAs such as tRNAs, as well as genes with incompletely characterized or mis-annotated ORFs. In addition, because they are not encoded entirely by successive triplets, it also eliminated two genes (*prfB* and *dnaX*) that contain +1 programmed frameshifts. The method selected 4119 ORFs containing ∼1.25 million codons. This large, essentially whole genome set should allow for robust, biologically relevant observations and correlations.

We make extensive comparisons between the second and third codons and their amino acids to those at more internal positions. We considered internal positions as those from the fourth codon through the penultimate sense codon. The last sense codon was not included to avoid possible bias related to termination efficiency^13^.

### How far does the bias at initiation sites extend before it transitions to internal bias?

The 5’ ends of coding sequences exhibit biases in nucleotide, codon and amino acid usage, and these biases differ from those of internal sequences. To determine the extent that start site biases into open reading frames, we examined their average nucleotide composition beginning with the fourth nucleotide (Figure 1A). ORFs begin with an overabundance of adenosine, which decreases and becomes accompanied by an increase in thymidine; and then by about the 20^th^ nucleotide, these patterns become replaced by a triplet G-T-nonA pattern that is phased with translation, and which is stable throughout coding sequences (not shown). The G-T-nonA pattern is similar to the G-nonG-N pattern recognized by Trifonov (1992) for a subset of coding *E. coli* sequences^8^. These results are consistent with A-richness and G-avoidance in the near vicinity of start codons being important to minimizing secondary structure that could occlude initiation sites. They also suggest that initiation-related sequence biases do not extend deeply into coding sequences, which would be consistent with models in which only a short region needs to be accessible for initiation.

**Figure 1.**
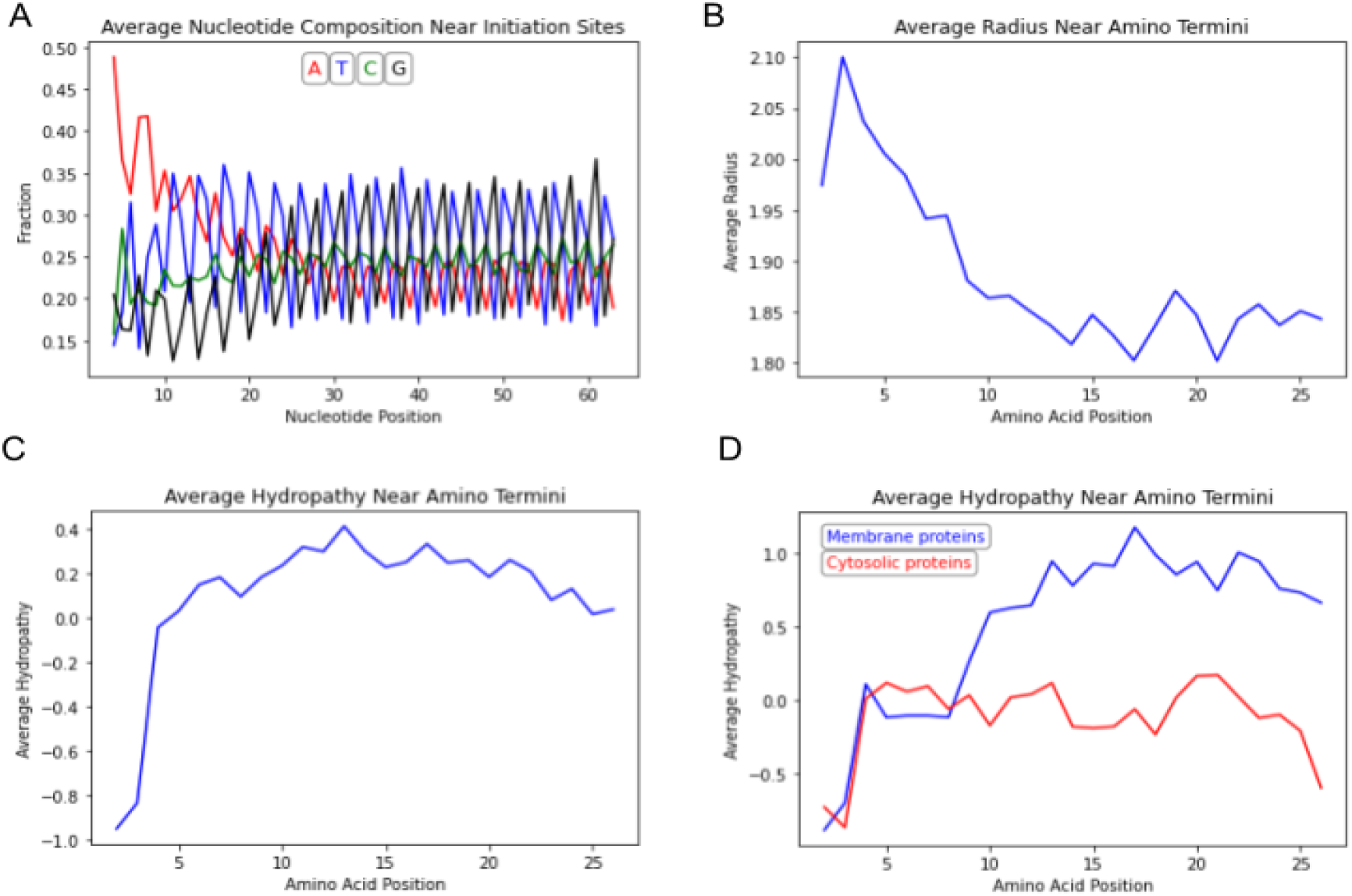
Characteristics of the 5’ ends of ORFs. (A)Average nucleotide composition by open reading frame position starting at the fourth nucleotide. (B) Average amino acid radius starting with the second position. (C, D) Average amino acid hydropathy starting with the second amino acid. (D) Presumptive integral membrane proteins (blue curve) were identified by keywords: ‘efflux’, ‘ABC’, ‘antiport’, ‘symport’, ‘permease’, ‘periplasm’, ‘ATP synthase’, ‘transport’, ‘quinone oxidoreductase’, ‘sensor’). Presumptive cytosolic protein genes (red curve) were identified by keywords: ‘ribosomal’, ‘ribosome’, ‘repressor’, ‘activator’, ‘polymerase’, ‘hydrolase’, ‘epimerase’, ‘isomerase’, ‘oxidoreductase’, ‘dehydrogenase’, ‘aldolase’, ‘hexokinase’, ‘mutase’, ‘enolase’, ‘kinase’, ‘citrate’. were identified by keywords: ‘ribosomal’, ‘ribosome’, ‘repressor’, ‘activator’, ‘polymerase’, ‘hydrolase’, ‘epimerase’, ‘isomerase’, ‘oxidoreductase’, ‘dehydrogenase’, ‘aldolase’, ‘hexokinase’, ‘mutase’, ‘enolase’, ‘kinase’, ‘citrate’.

Since the amino terminal regions of proteins can have important roles in protein processing, we also wished to determine whether the amino acid nearest neighbors of the universal f-Met would be biased for size and/or hydrophobicity. Figure 1B shows that the average size of amino acids 2 through 9 = 1.98 Å, while the average from amino acids 12 to 19 = 1.83 Å. This difference could be due in part to the overabundance of lysine, and paucities of valine, alanine and glycine, at the beginnings of ORFs (addressed below).

To examine average hydrophobicity, we used the Kyte and Doolittle (1982) scale of “hydropathy” (relative hydrophobicity estimated by combining several types of measurement) in which hydrophobic amino acids have positive scores (up to 4.5 for isoleucine) and hydrophilic amino acids have negative values (down to −4.5 for arginine)^14^. We plotted the average hydropathies of the amino acids near the amino termini of all proteins in figure 1C, which shows that amino acids 2 and 3 have relatively low average hydropathies and the remainder have average hydropathies of ∼0.2. We also plotted the data for two subset of genes that should differ in their overall average hydropathies (Figure 1D): genes for integral membrane or cytosolic proteins (531 and 611 genes, respectively; see legend of fig. 2 for details). Figure 1D shows that beginning at about the tenth amino acid, integral membrane proteins have a higher average hydropathy internally than cytosolic proteins do, which should not be surprising (reviewed by White and Wimley, 1999)^15^. However, the two protein sets have similar, and low, hydropathies at their second and third amino acids.

**Figure 2.**
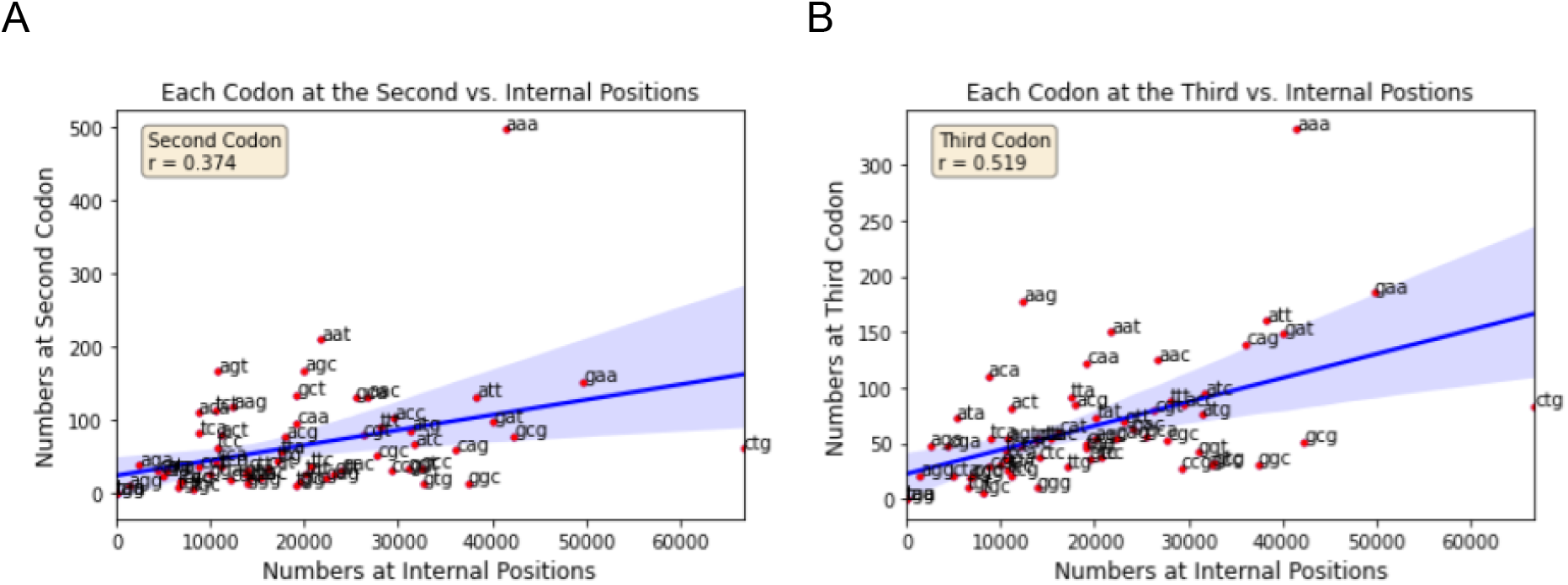
Codon bias near initiation sites. The numbers of each sense codon internally plotted vs. the second (A) and third (B) positions. Linear correlation coefficients (r) are shown in the insets, and shaded areas define the 95% confidence intervals.

To summarize the results in Figure 1, there is strong bias for adenine beginning at the 4^th^ nucleotide, which gradually fades by about the 20^th^ nucleotide. As adenine declines, thymidine increases; and then both A- and T-richness are replaced by an extended G-T-nonA internal pattern. The initiation site-proximal biases are associated with the usage of amino acids whose average physical properties differ from the subsequent internal amino acids. These bases are strongest for the second and third amino acids. We now address the possible functional significance of these observations.

### What are the likely causes of initiation region bias?

#### A-richness is a major but not the only factor driving ORF initiation site-proximal bias

To compare codon bias near initiation to that of internal sites, we plotted the numbers of each sense codon at the second or third positions vs. those at internal positions (Figure 2). The correlations are very poor, and especially notable is that AAA is strongly favored as both the second and third codons. Further inspection shows that codons above the best fit lines tend to be A- and/or U-rich. Together, these observations are consistent with the hypothesis that initiation sites are under selection to lessen the potential for secondary structure to provide for access by ribosomes^16,17,18^. In addition, at the second position the AGT and AGC serine codons plot well above the best-fit line and are thus overrepresented there. In contrast, they are not overrepresented at the third position: AGC is just below the line near coordinates [28000, 50], and AGT is not clearly visible but is a cluster of codons just below the line near coordinates [20000, 50]. Below, we return to a possible special role for second-position serine in f-Met removal.

There is, however, considerable complexity in these plots that may be caused by factors in addition to A-richness. To explore this complexity further, we plotted observed codon frequencies versus their frequencies expected by assuming that triplet probabilities are the products of their constituent nucleotide frequencies (Figure 3). For example, the expected frequency of AAA at the second codon is the product of frequencies of adenosine at nucleotides 4, 5 and 6. The correlation coefficients (r values) are substantially higher than those in figure 2, which suggests that codon bias at these positions is strongly dependent on nucleotide bias. Although correlation does not prove causation, the coefficients of determination (r^2^) between observed and expected codon frequencies for both the second and third codons are about 54% and 70%, respectively, which suggests that roughly ½ to ¾ of codon bias could be explained by nucleotide bias. The remainders may have other causes that we return to below.

**Figure 3.**
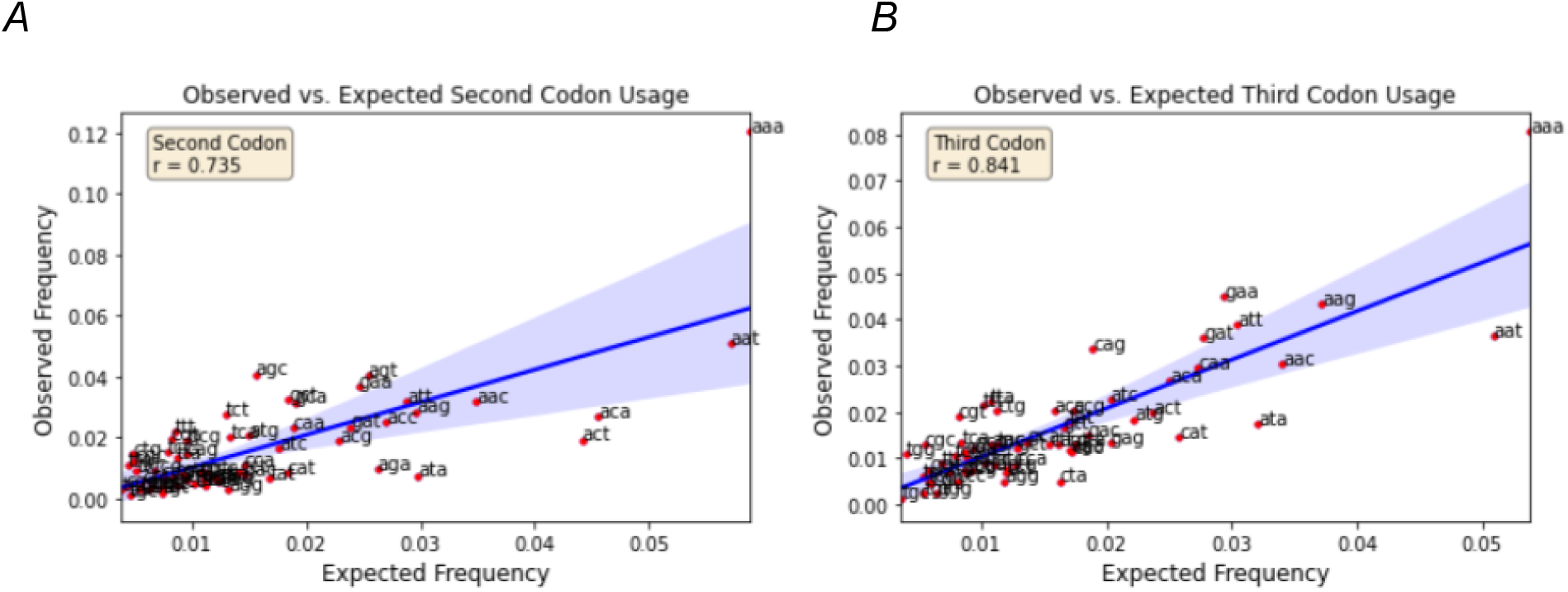
Sense codon frequencies vs. expected frequencies. Frequencies expected from nucleotide frequencies for the second (A) and third (B) positions. Linear correlation coefficients (r) are shown in the insets, and shaded areas define the 95% confidence intervals.

One feature that stands out in all of the figure 2 and 3 plots is the overabundance of the AAA triplet. It seems likely that either AAA or lysine is important for some function in at least a large fraction of genes. To determine whether or not the overrepresentation of triple-A is strictly due to its coding function, we determined whether triple-A is overrepresented in the two non-translational reading frames between the second and third in-phase codons (Figure 4).

**Figure 4.**
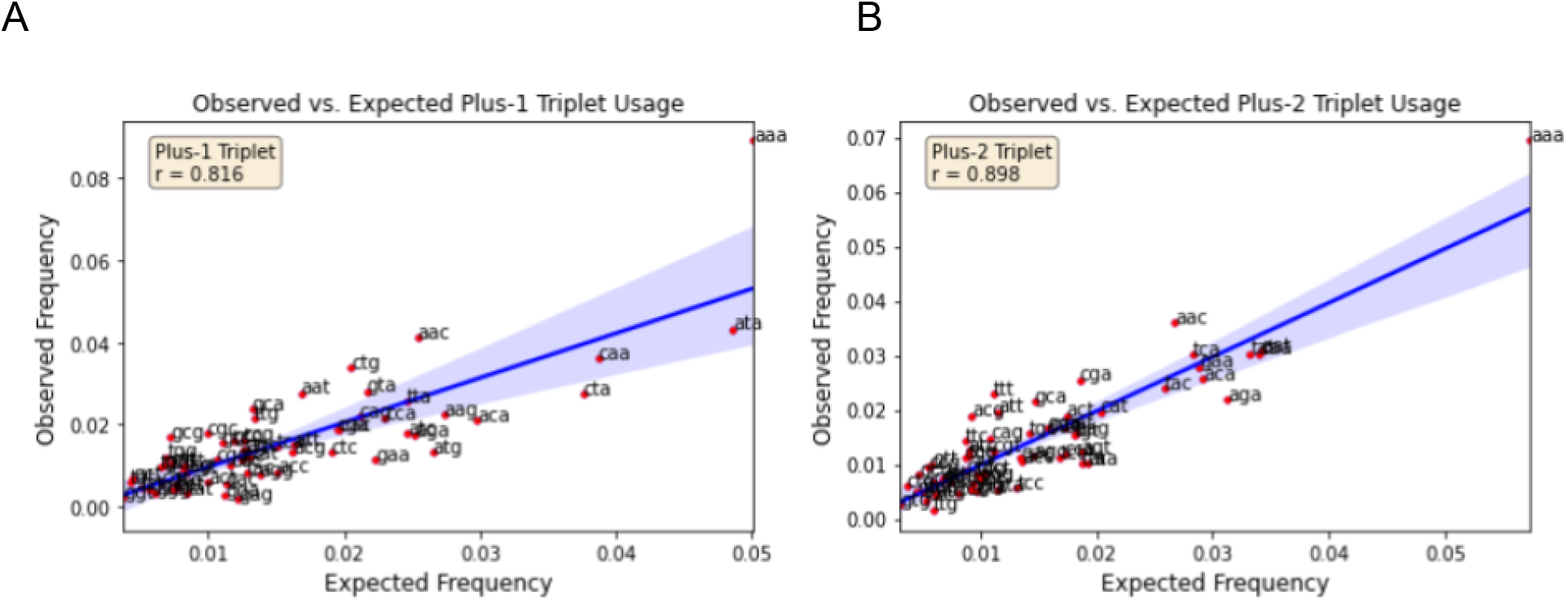
Observed vs. expected frequencies of the triplets located in the two phases between the second and third codons. The plus-1 triplet is nucleotides 5, 6 and 7. The plus-2 triplet is nucleotides 6, 7 and 8. Linear correlation coefficients (r) are shown in the insets, and shaded areas define the 95% confidence intervals.

It is clear from figures 3 and 4 that the AAA triplet is overrepresented in every reading frame including the non-translated frames. These results strongly suggest that triple-A is selected at least partly for a noncoding function. For example, three consecutive adenosines are almost certainly more effective at disrupting secondary structures than would be expected from the effects of individual adenosines, which can be accommodated as mispairs or bulges in RNA structures (RNA structural analysis has been recently reviewed by Li et al, 2020)^19^.

Although nucleotide bias, especially for A-richness, is likely to be a substantial cause of codon choice, it is not the only cause. Please note that A-richness does not decrease smoothly into ORFs. The plot for individual nucleotide frequencies (Figure 1) shows that adenine frequency decreases in a sawtooth rather than a smooth pattern. Such unevenness suggests that other pressures also affect sequences, which should not be surprising. Such effects may be complex, and perhaps idiosyncratic to specific sets of genes or proteins. For example, highly and weakly expressed genes are well-known to exhibit different patterns of highly nonrandom codon and context usage, the mechanisms for which are incompletely understood^18,19^. We did not include expression level or other factors as variables in these analyses. But to detect potential, broadly applicable causes of these initiation sequence biases, we examined the frequencies of the second and third amino acids.

#### Bias involving the second and third amino acids

Previous work on a subset of genes (Looman et al. 1987) or of the Swiss-Prot protein database (Shemesh et al. 2010) had identified amino acid biases at the N-termini of *E. coli* proteins^9,22^. We extended that work with an examination of the ORFs from the entire *E. coli* K12 genome. These results are shown in Figure 5. There, the 20 amino acids are rank-ordered, left-to-right, by their frequencies at internal positions (e.g., Leucine > Alanine > Glycine …). There are three colored bars for each amino acid: blue for internal, red for the second, and green for the third amino acid positions.

**Figure 5.**
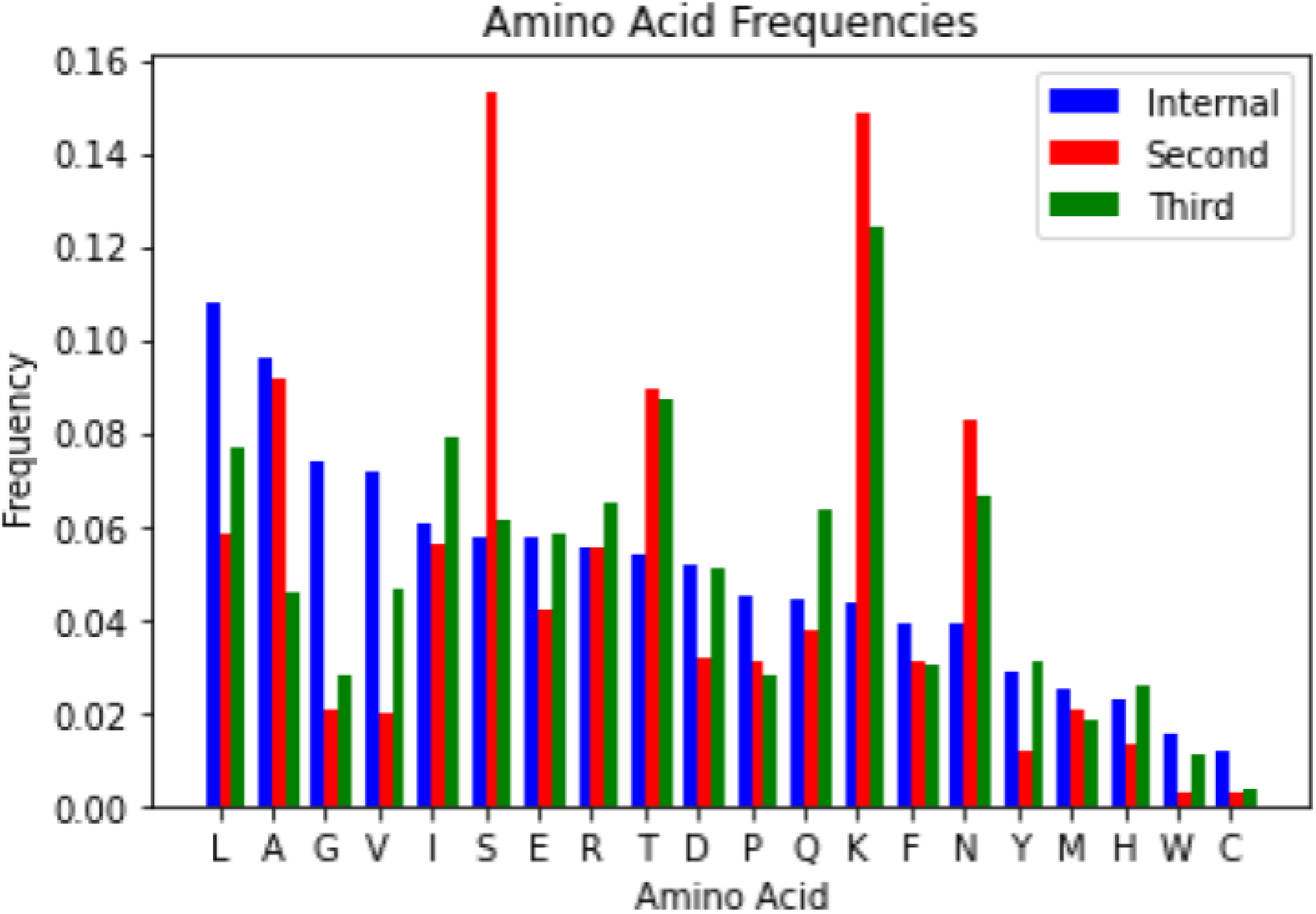
Amino acid bias near amino-termini. Amino acids are rank-ordered by the frequencies at internal positions (blue bars). To the right of each bue bar, the frequency of the corresponding amino acid at the second position is shown as a red bar, and at the third codon as a green bar.

Figure 5 shows that the serine, lysine, threonine and asparagine are strongly over-represented relative to their usage internally. Together, these four hydrophilic amino acids comprise about 45% and 35% of the second and third amino acids, respectively. In contrast, most of the hydrophobic amino acids that are very common internally such as leucine, alanine and valine are underrepresented as the second amino acid. One consequence of these patterns is that average hydropathy of the second and third amino acids is relatively low (see Figure 1 panels C & D), which could have functional significance.

It is possible that these near-terminal amino acid biases could at least partly be due to effects on protein processing. Most *E. coli* proteins have their terminal f-Met removed, possibly as a signal to control protein half-life^23^. Moreover, methionine is a metabolically expensive amino acid (reviewed in Sekowska et al. 2019)^24^, and its efficient removal for reuse would be energetically economical. Its removal by the successive action of two enzymes, peptide deformylase and methionyl aminopeptidase, is sensitive to the second amino acid^23^. However, second amino acid frequency bias does not correlate in a simple way with the known effects on f-Met removal^9^. For example, removal is facilitated if the second amino acid is relatively small and moderately polar. If second amino acid bias were strictly due to effects on f-Met removal one might expect that all such amino acids would exhibit similar usage biases; but that is not the case. Consider, for example, that second position glycine, alanine, serine, cystine, proline, threonine, and valine all serve as good substrates but only serine and threonine are overrepresented in *E. coli* genes^25^. Similarly, lysine and leucine would make poor substrates, but lysine is highly favored and leucine is avoided. One feature that unites all of the favored second amino acids is the possibility of specification by A- or A,U-rich codons. Thus it seems possible that two factors contribute to apparent amino acid bias, initiation site openness and f-Met removal, and these factors may sometimes be in conflict. Consider that lysine is strongly overrepresented at second position despite being a poor substrate for f-Met removal. But because it is specified by the AAA triplet, lysine’s abundance is likely due to the selection for initiation site availability as discussed above.

Several observations support a functional role for serine as the second amino acid, and facilitation of f-Met removal is a likely function. Also, because it can be specified by codons that contain adenosine and thymidine, it may be able to serve one or more functions without sacrificing initiation site openness. Intriguingly, serine is the most common second amino acid (∼15%), but it is not overrepresented as a third amino acid (figure 5). Moreover, the serine codons AGT and AGC are strongly overrepresented as the second codon both relative to internal usage (figure 2A), and to usages expected from nucleotide frequencies (figure 3A). However, none of the serine codons are overrepresented at the third (figures 2B and 3B) or at plus-1 or plus-2 positions (figures 4). The overrepresentation only at the second position strongly suggests that it is useful as a second amino acid.

To address this issue more fully, we examined the usage of the six serine codons (Figure 6). Overall, serine codons are 2.5-times more common at the second position than they are at internal positions (Figure 6), and the increases are highly significant. Codons whose increase favors A/U-richness show greater relative increases than others. For example, AGT and AGC are read by the same tRNA, but while AGC is preferred internally, they are used equally at the second codon. Similarly, TCT and TCA both have increased usage at the second position, while TCC barely does, and TCG does not increase at all. Together, these observations suggest that second position serine is somehow favorable to proteins or to protein synthesis, possibly because it facilitates f-Met removal, and that this is done through increases of serine codons that promote A-richness, thus helping to keep initiation sites accessible to initiating ribosomes.

**Figure 6.**
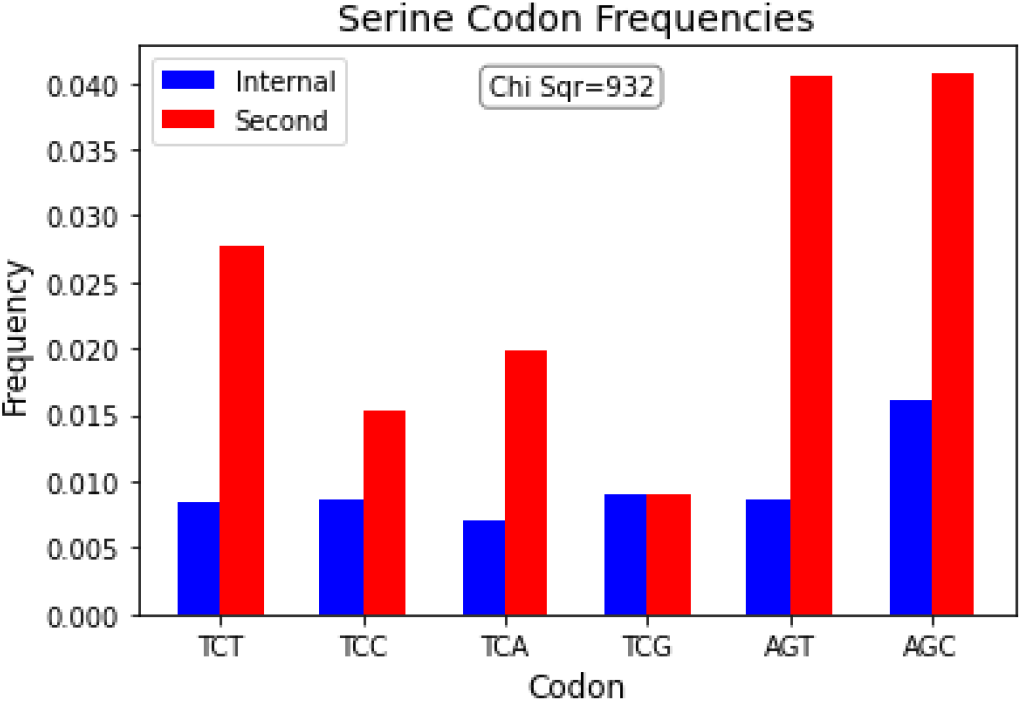
Usage of the serine codons internally and at the second position. χ^2^ value of 932 has a probability <<0.001 (five degrees of freedom).

We also considered whether the second and third amino acids might facilitate protein secretion. There are several pathways for protein translocation into or across the plasma membrane that can depend on the physical properties of the amino acids of ∼5-30 amino acid secretory signal peptides (SPs), usually located near the amino terminus^26,27^ (recently reviewed by Owji et al, 2018 and Hui et al., 2021)^28,29^. But the details are too complex to be unraveled by our analysis of average features. Nonetheless, because the N-proximal regions of SPs often contain basic amino acids, it is possible that the overabundance of lysine as the second and third amino acids (figure 5) facilitates the export of at least some proteins. However, because the other strongly basic amino acid, arginine, is not overrepresented there, selection for export is unlikely to be a significant driver of amino acid bias for the second and third amino acids. Instead, the abundance of lysine is likely due primarily to the preference for AAA triplets discussed above.

Other plausible roles for the amino terminal amino acids include entrance, passage and exit of the nascent peptide through the ribosomal polypeptide exit tunnel. Although the long, hydrophobic f-Met side chain should enhance entrance into the tunnel (see discussion in Lim et al., 2012)^30^, its formylated amino group and hydrophobic side chain could render the f-Met terminus slow to solvate at the end of the tunnel. Therefore, the relatively low average hydropathy for the second and third amino acids (Figures 2 C & D), due to the abundance of serine, lysine, threonine and asparagine (Figure 6) could facilitate exit of the amino terminus into the cytoplasm by promoting its solvation.

In conclusion, nucleotide, codon and amino acid biases may have many causes, which may sometimes conflict. But A-richness, presumably to maintain initiation region accessibility, is a principal one. The facilitation of f-Met removal, especially by serine, is also a likely important force. It is also possible that low hydropathies for the second and third amino acids facilitate polypeptide emergence from the ribosomal exit channel.

### Does bias at initiation sites differ among genes that use AUG vs. other initiation codons?

Approximately 10% of *E. coli* genes have non-AUG start codons. Because translation initiation always uses the same tRNA, base pairing at non-AUG initiation codons must involve wobble pairing at one or more positions. At least formally, then, initiation at non-AUG sites may be prone to problems related to suboptimal base pairing such as decreased accuracy. We reasoned that any such problems might create selective pressures on the second or third codons to somehow compensate for them. Therefore, we compared codon usage at these codons between AUG and non-AUG starting genes (Figure 7). The very strong correlations suggest that any difficulties associated with initiation with non-AUG codons, if they exist, do not impose strong selective pressures on second and third codon identity. One possible exception might be that the A-rich asparagine codons (AAT, AAC) are increased at the second but not third codon for non-AUG starting genes. Like serine, asparagine should facilitate f-Met removal, but its codons further promote A-richness, perhaps to compensate for suboptimal initiator codons. This signal is relatively small, however, involving only about 10 genes. The nucleotide compositions at the second and third codon positions are not distinguishable between AUG- and non-AUG-starting genes (not shown).

**Figure 7.**
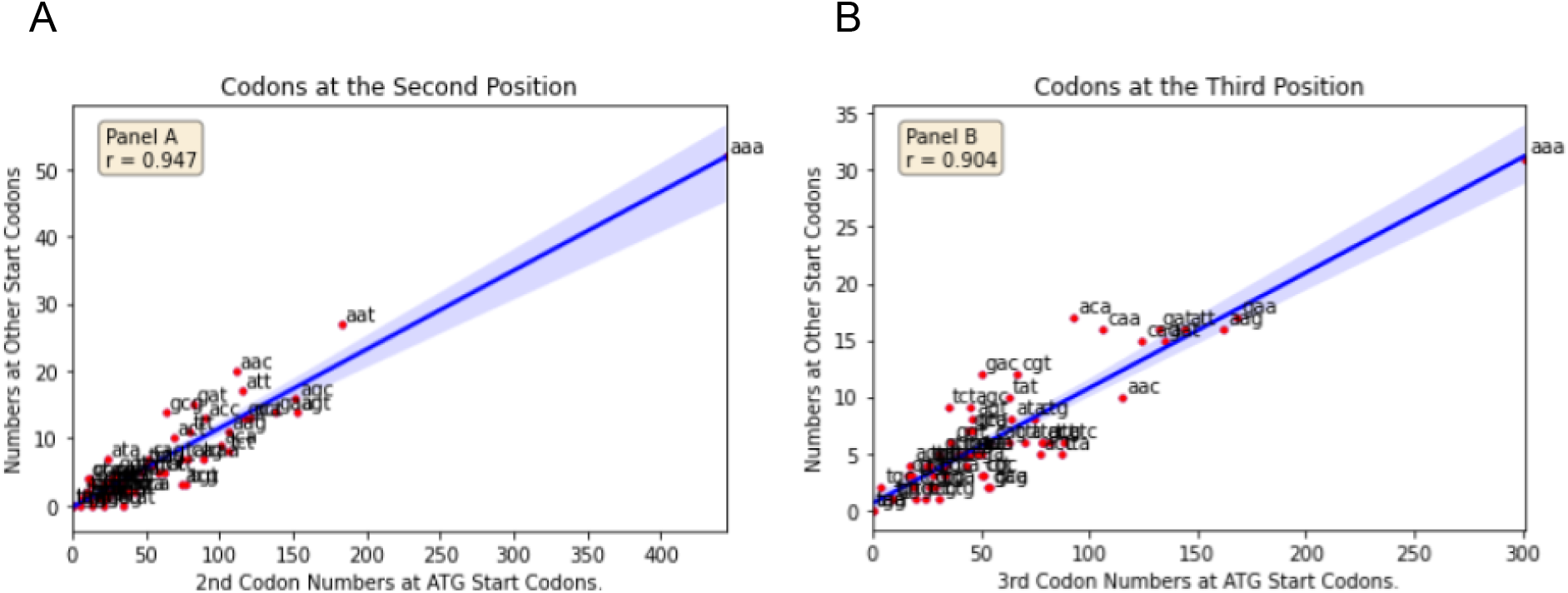
Codon Usage Near Start Codons. Usage of the second (A) and third (B) codons of gene that begin with AUG vs. other initiation codons. Linear correlation coefficients (r) are shown in the insets, and shaded areas define the 95% confidence intervals.

### Does initiation codon identity affect the establishment of the correct reading frame?

We hypothesized that if mRNA:tRNA duplex stability during initiation is functionally significant in establishing the reading frame, then genes that initiate with non-AUG codons may be relatively frameshift prone. To test this idea, we created constructs that included features of the *prfB* programmed frameshift site, for which frameshift frequency has been observed to vary 17-fold depending on the stability of *cognate* mRNA:tRNA duplexes in the E-site^31^. Our constructs were created as described in the Methods, and the frameshift-prone site is outlined in Figure 8. During the frameshift, the P site tRNA slips by one nucleotide into the +1 reading frame (as indicated by the red bars in the figure) translation then continues in that new frame. Frameshifting is enhanced when the A site codon is slowly translated^32,33^. In our constructs, the A site contains CGG, which is a rare and slowly translated codon in *E. col*i^33^. Critically for this experiment, the frequency of such slippage is expected to be sensitive to the stability of the initiation codon (NUG):initiator-tRNA duplex, now in the E-site, such that unstable duplexes should increase the probability of the frameshift.

**Figure 8.**
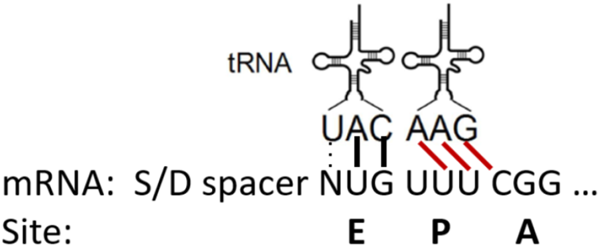
The translational complex at the time of the frameshift. The initiation codon (NUG) is in the E-site. The P-site tRNA can slip from the in-phase UUU onto the UUC triplet in the +1 frame (red bars indicate base pairs after slippage).

We compared *lacZ* expression between four pairs of constructs. The pairs differed in their initiation codons (AUG, GUG, CUG, UUG), and each pair consisted of alleles that did and did not require frameshifting for *lacZ* reporter expression (Methods). Alleles that do not require frameshifting (in-frame alleles) serve as controls for translational initiation efficiency, which is known to differ among these four initiation codons^34^. The data (Table 1) show that the in-frame AUG construct has an activity similar to that of the parent plasmid control, which also has an AUG initiation codon. Among the NUG in-frame set, initiation efficiency varies in the order AUG > GUG > UUG > CUG.

**Table 1.** Frameshifting frequencies of the frameshift prone (UUU_C) sequence.

| Test Sequence | Codon-Anti | Activity | %Normal |
| --- | --- | --- | --- |
| pJC1168 | UAC<br>AUG | 28546.512 |  |
| AUG UUU CGG AAC | UAC<br>AUG | 29075.464 |  |
| AUG UUU CGG U AAC | UAC<br>AUG | 35.958 | 0.001 |
| CUG UUU CGG AAC | UAC<br>CUG | 718.303 |  |
| CUG UUU CGG U AAC | UAC<br>CUG | 1.529 | 0.002 |
| UUG UUU CGG AAC | UAC<br>UUG | 1005.603 |  |
| UUG UUU CGG U AAC | UAC<br>UUG | 1.048 | 0.001 |
| GUG UUU CGG AAC | UAC<br>GUG | 4508.944 |  |
| GUG UUU CGG U AAC | UAC<br>GUG | 4.98 | 0.001 |

All of the frameshift alleles exhibit readily measurable frameshift-dependent activity, which shows that the assay is a sensitive probe of the frameshift mechanism. Importantly, however, the “% Normal” frameshift activities indicate that frameshift events occur at frequencies of only about 1 and 2 per thousand, which are comparable to background values from other experimental systems. Importantly, there is no significant increase for any of the NUG variants. These observations suggest that the quality of the start codon:tRNA interaction in the E-site does not affect the probability that the tRNA will prematurely leave the E-site, which might have interfered with the establishment of the reading frame.

## Conclusions

The beginning of the average *E. coli* ORF is highly non-random, and patterns of nucleotide, codon and amino acid usage are very different from those of average internal sequences. Multiple mechanisms are likely to drive these biases, and although it is likely that many genes have their individual reasons, there are several likely general causes. For example, A-richness, which can help maintain initiation region accessibility, is a principal one^17,35^. Moreover, the AAA triplet is overrepresented in all reading frames, suggesting that it is important beyond its coding function. It is highly likely that three consecutive adenines is very effective at preventing message structures that would otherwise include initiation sites. The facilitation of f-Met removal by several small, moderately hydrophilic amino acids is also a likely important driving force. Serine may be particularly important as a second amino acid because its use can promote both message A,U-richness and f-Met removal. The possibility that low hydropathies of the second and third amino acids promote protein transit or exit from the polypeptide exit tunnel is intriguing.

About 10% of *E. coli* genes have non-AUG initiation codons, which are known to have reduced initiation efficiencies, suggesting that their codon:anticodon interactions may be relatively weak^34^. However, we did not detect substantial differences in the usages of second and third codons between AUG- and nonAUG-starting genes. We also did not detect increased frameshifting due to tRNA dissociation from the ribosomal E-site. Thus, weak pairing between the initiator tRNA and non-AUG initiation codons may not impair subsequent translational events.

## Acknowledgements

We are grateful to Dr. David Ornelles for comments on the manuscript. This work was supported by NIH Grant GM-077194 to J.F.C.

## Notes

https://github.com/Curran-WFU

### Competing Interest Statement

The authors have declared no competing interest.

